# Intranasal administration of a VLP-based vaccine against COVID-19 induces neutralizing antibodies against SARS-CoV-2 and Variants of Concerns

**DOI:** 10.1101/2021.09.10.459749

**Authors:** Dominik A. Rothen, Pascal S. Krenger, Aleksandra Nonic, Ina Balke, Anne-Cathrine S. Vogt, Xinyue Chang, Alessandro Manenti, Fabio Vedovi, Gunta Resevica, Senta M. Walton, Andris Zeltins, Emanuele Montomoli, Monique Vogel, Martin F. Bachmann, Mona O. Mohsen

## Abstract

**Background:** The highly contagious SARS-CoV-2 is mainly transmitted by respiratory droplets and aerosols. Consequently, people are required to wear masks and maintain a social distance to avoid spreading of the virus. Despite the success of the commercially available vaccines, the virus is still uncontained globally. Given the tropism of SARS-CoV-2, a mucosal immune reaction would help to reduce viral shedding and transmission locally. Only seven out of hundreds of ongoing clinical trials are testing the intranasal delivery of COVID-19 vaccines.

**Methods:** In the current study, we tested in murine model the immunogenicity of a conventional vaccine platform based on virus-like particles (VLPs) displaying RBD of SARS-CoV-2 for intranasal vaccination. The candidate vaccine, CuMV_TT_-RBD, has been immunologically optimized to incorporate tetanus-toxin and is self-adjuvanted with TLR7/8 ligands.

**Results:** CuMV_TT_-RBD elicited strong RBD- and spike- specific systemic IgG and IgA antibody responses of high avidity. Local immune responses were assessed and results demonstrate strong mucosal antibody and plasma cell production in lung tissue. The induced systemic antibodies could efficiently recognize and neutralize different Variants of Concerns of mutated SARS-CoV-2 RBDs.

**Conclusion:** In summary, intranasal vaccination with CuMV_TT_-RBD shows high immunogenicity and induces protective systemic and local specific antibody response against SARS-CoV-2 and its variants.

**One sentence summary:** Evaluation of an intransal administrated conventional VLP-based vaccine against COVID-19 in a murine model.

## 1. Introduction

Up to date, COVID-19 caused by SARS-CoV-2 is still considered a global pandemic that has wreaked havoc globally and put a heavy toll on public health and economy. The marketed vaccines such as mRNA, viral vector and inactivated viruses have greatly reduced the number of COVID-19 mortality and hospitalization and continue to provide different levels of protections against the emerged and emerging Variants of Concerns (VOCs)^1^.

Viral tropism depends, amongst other factors, on the susceptibility of a specific host cell. COVID-19 patients often present with respiratory illness that can progress to severe pneumonia^2^. Such observations suggested that the lung is the primary organ infected by SARS-CoV-2, which is consistent with angiotensin converting enzyme 2 (ACE2), which expressed by lung epithelial cells, as the viral receptor ^3^. The primary port of entry to the body are alveolar epithelial cells, but vascular endothelial cells also express ACE2 and are a prominent place of viral replication^4,5^. These cells may be considered the base for early infection and viral replication as well as long-term viral persistence in some cases^6^.

The currently available marketed vaccines are administered intramuscularly (i.m.) producing systemic spike and RBD-specific antibodies (Abs) that can recognize and neutralizes the virus^7^. Given the tropism of SARS-CoV-2, recent research efforts have also been devoted towards the development of an intranasal COVID-19 vaccine. Seven intranasal COVID-19 vaccines candidates are in clinical trials^8^. Intranasal (i.n.) vaccination route may offer several advantages over i.m. route including: needle-free administration, direct delivery to the site of infection and most importantly the induction of mucosal immunity in the respiratory tract ^9^. Secretory IgA (sIgA) is of major importance in the respiratory tract where it presents an efficient line of defence against respiratory infections^10^. Furthermore, mucosal vaccination can result in resident B and T cell priming leading to long-lived Ab secreting cells or tissue-resident memory cells, which add in clearing the viral infection^11^. This locally induced immune reaction has been shown to reduce viral replication and shedding in lungs and nasal passages leading to lower infection and transmission^12^. The concept of i.n. vaccination goes back to 1960s based on observations with live-attenuated influenza vaccines (LAIV) that mimic a natural influenza infection and have shown to elicit protective a local and systemic antibody as well as cellular responses^13^. Live-attenuated virus or viral-vector based vaccines need to infect cells for replication. Also, attenuated viruses may pose a small risk of retaining their replication ability, especially in people with weaker immune systems.

Efficient induction of mucosal immunity can be best achieved by vaccines that mimic mucosal pathogens. Virus-like particles (VLPs) constitute an efficient and safe vaccine platform as they lack genetic materials for replication *in vivo*. Their particulate repetitive surface structure enables them to stimulate innate and adaptive immune response and target the mucosa as well as the underlying dendritic cells (DCs)^14^. We have previously assessed the efficacy of Qß-VLPs as an i.n. vaccine platform. Our results indicated efficient induction of specific-IgGs in serum and lung besides robust local IgA production^14^. Here we provide a proof-of-concept (PoC) in murine model proving the immunogenicity and efficacy of i.n. administration of a SARS-CoV-2 vaccine based on VLPs. The developed vaccine candidate is based on our optimized plant-derived VLPs (CuMV_TT_-VLPs) displaying the receptor-binding domain (RBD) of SARS-CoV-2. Our results demonstrate that CuMV_TT_-RBD induces strong systemic and local B cell response including high levels of IgG and IgA, plasma cell (PC) formation as well as broad viral neutralization. Taken together, our vaccine constitutes an efficient candidate for the generation of Ab-based vaccine that can be administered mucosal in a needle-free manner.

## 2. Results

### 3.1 CuMV_TT_ VLPs constitute an efficient platform for vaccine development

In order to generate a vaccine-candidate against SARS-CoV-2 for i.n. administration, we have utilized our optimized plant-derived VLPs (CuMV_TT_-VLPs) as a vaccine platform^19,15^. RBD amino acid sequence (a.a. Arg319-Phe541) of SARS-CoV-2 was chemically coupled to CuMV_TT_-VLPs using SMPH bifunctional cross-linker (Fig. 1a). The generated vaccine candidate CuMV_TT_-RBD is self-adjuvated with prokaryotic ssRNA (TLR7/8 agonist) which is packaged during expression and assembly in the bacterial *E. coli* system (Fig. 1b). Efficiency of RBD coupling to CuMV_TT_-VLPs was confirmed by SDS-PAGE (Fig. 1c). The integrity of the VLPs following the coupling process was checked by electron microscopy and showed no signs of aggregation (Fig. 1d).

**Figure 1.**
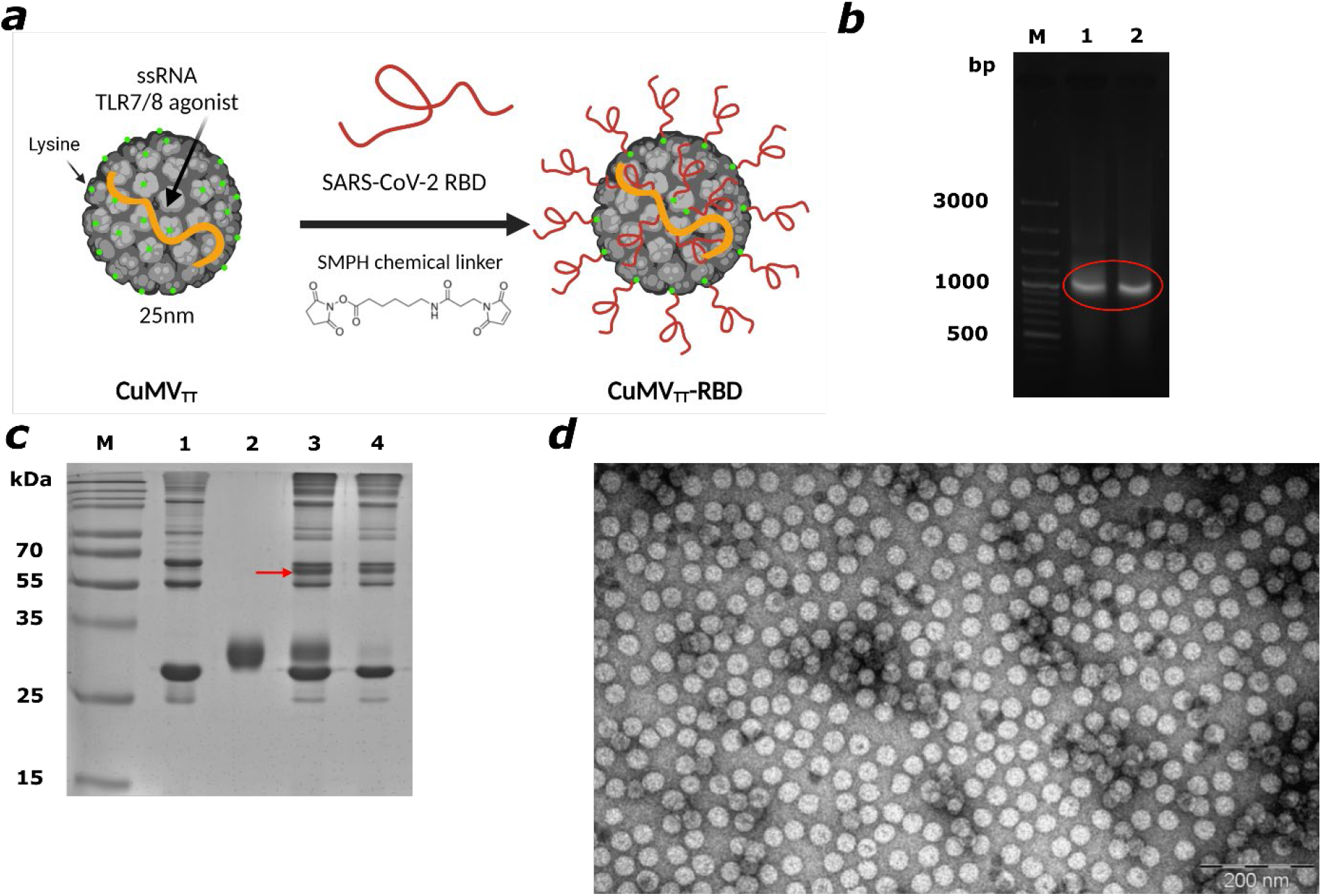
CuMV_TT_ VLPs constitute an efficient platform for vaccine development. a) Schematic representation of the chemical coupling of RBD to CuMV_TT_-VLPs via SMPH bifunctional cross-linker. b) Agarose gel analysis of CuMV_TT_-RBD and CuMV_TT_ depicting nucleic acids packaged in CuMV_TT_. M, DNA Ladder, 1. CuMV_TT_-RBD, 2. CuMV_TT_ VLP containing bands are labelled with red circle. c) 12% SDS-PAGE for CuMV_TT_-RBD production. M. Protein marker, 1. CuMV_TT_, 2. RBD, 3. CuMV_TT_-RBD pre-wash, 4. CuMV_TT_-RBD post-wash. Red arrow indicating the coupled CuMV_TT_-RBD product. d) Electron microscopy of CuMV_TT_-RBD, scale bar 200nm.

### 3.2 Intranasal administration of CuMV_TT_-RBD induces a systemic RBD- and spike-specific IgG response of high avidity

To test the immunogenicity and the induction of a humoral immune response in murine models, BALB/c mice were i.n. primed on day 0 and boosted on day 28 with 40μg of CuMV_TT_-RBD vaccine or with 40μg of CuMV_TT_-VLPs as a control without addition of adjuvants. Vaccination and bleeding regimen are shown in Figure 2a. Total systemic RBD and spike-specific IgG were measured by ELISA. Systemic RBD-specific IgG response was detected in the group received CuMV_TT_-RBD seven days after the priming dose. Furthermore, the induced response increased by about 1000-fold following the booster dose on day 35 (Fig. 2b and c). Full-length spike protein responses remained low after priming but increased significantly by 30-folds upon a booster injection resulting in a stable IgG antibody titer (Fig. 2d and e). No IgG response has been detected in the control group which received CuMV_TT_-VLPs only.

**Figure 2.**
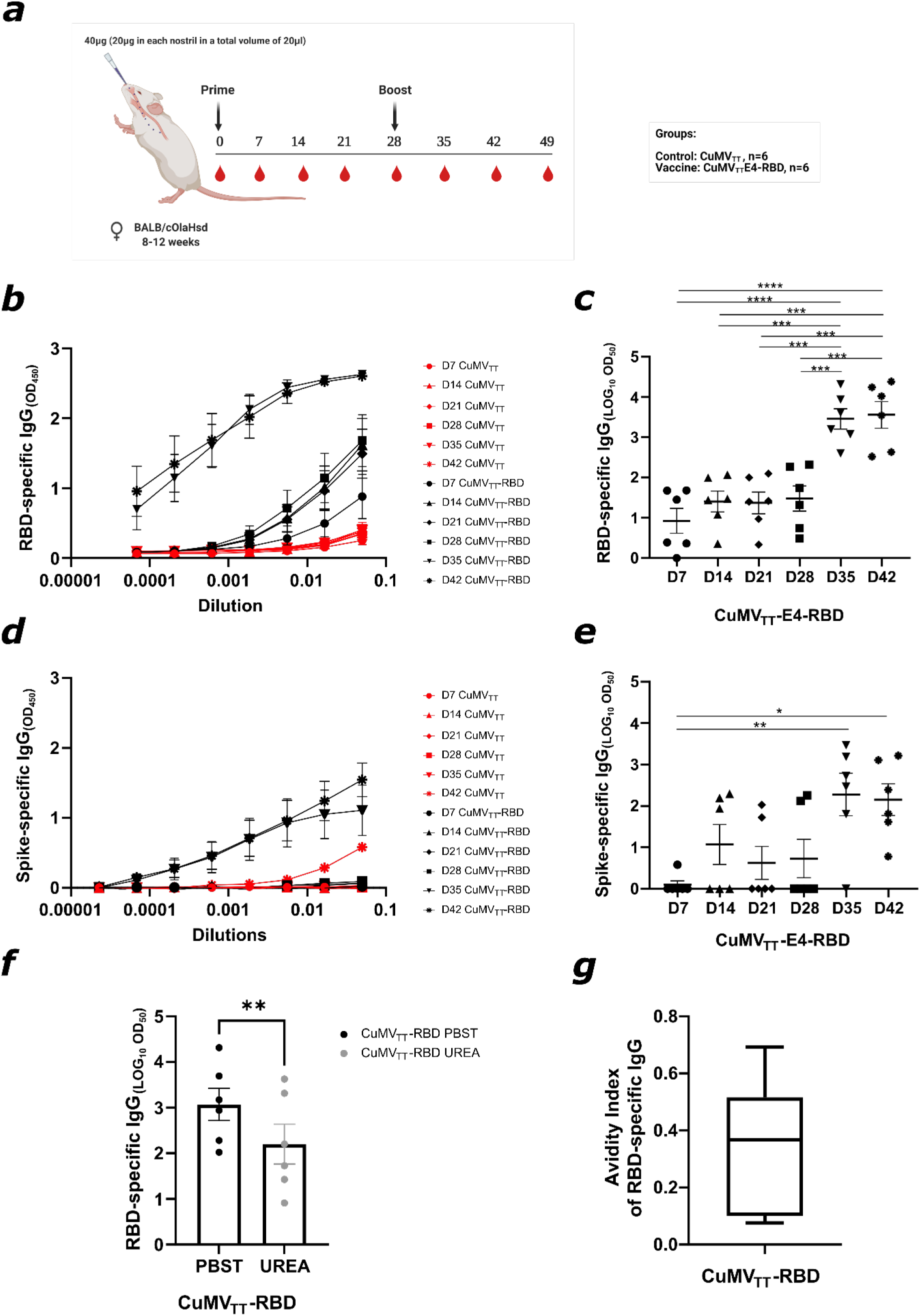
Intranasal administration of CuMV_TT_-RBD induces a systemic RBD- and spike-specific IgG response of high avidity. a) Vaccination regimen and bleeding schedule. b,c) RBD-specific IgG titer on days 7, 14, 21, 35 and 42 from mice immunized with CuMV_TT_-RBD vaccine or CuMV_TT_ control measured by ELISA, OD450 shown in b), LOG_10_ OD_50_ shown in c). d,e) Spike-specific IgG titer on days 7, 14, 21, 35 and 42 from mice immunized with CuMV_TT_-RBD vaccine or CuMV_TT_ control measured by ELISA, OD450 shown in d), LOG_10_ OD_50_ shown in e). f) RBD-specific IgG titer at day 49 from mice immunized with CuMV_TT_-RBD vaccine, LOG_10_ OD_50_ shown. g) Avidity index. Statistical analysis (mean ± SEM) using one-way ANOVA (c and e) and *Studen’t t-test* (f). Control group *n=6*, vaccine group *n=6*. One representative of 2 similar experiments is shown.

The avidity of an Ab is defined as the binding strength through points of interaction. It can be quantified as the ratio of *K*_d_ for the intrinsic affinity over the one for functional affinity of a single point interaction^19^. High avidity Abs are formed upon affinity maturation in germinal centers (GCs) and are associated with protective immunity against SARS-CoV-2 infection^20^. To assess the avidity of the induced IgG Abs against RBD, we carried out an avidity ELISA using day 49 sera. The obtained results indicated that about 40% of the systemically induced RBD-specific IgGs are of high avidity following the i.n. vaccination (Fig. 2 f and g).

### 3.3 Intranasal immunization with CuMV_TT_-RBD promotes isotype switching to IgA and leads to balanced IgG subclass responses

IgG subclasses are of major importance in the immunological response against viruses because of enhancing opsonization as well as immune effector funcitons^21^. Additionally, IgA plays an important role in protection against respiratory viruses as it is found in mucosal tissue, the main entrance site for these kind of viruses^10^. The ability of the CuMV_TT_-RBD vaccine to induce serum IgA and IgG subclasses was evaluated by performing ELISA against RBD with sera collected at day 42. All IgG subclasses were induced in vaccinated mice with IgG1, IgG2a and IgG2b being the dominant ones. In contrast, no IgG subclasses were detected in the control group (Fig. 3a). The vaccine was also able to induce isotype switching to IgA as shown in Fig. 3b. Approximately, 20% of serum IgA Abs were of high avidity as shown in (Fig. 3c-d).

**Figure 3.**
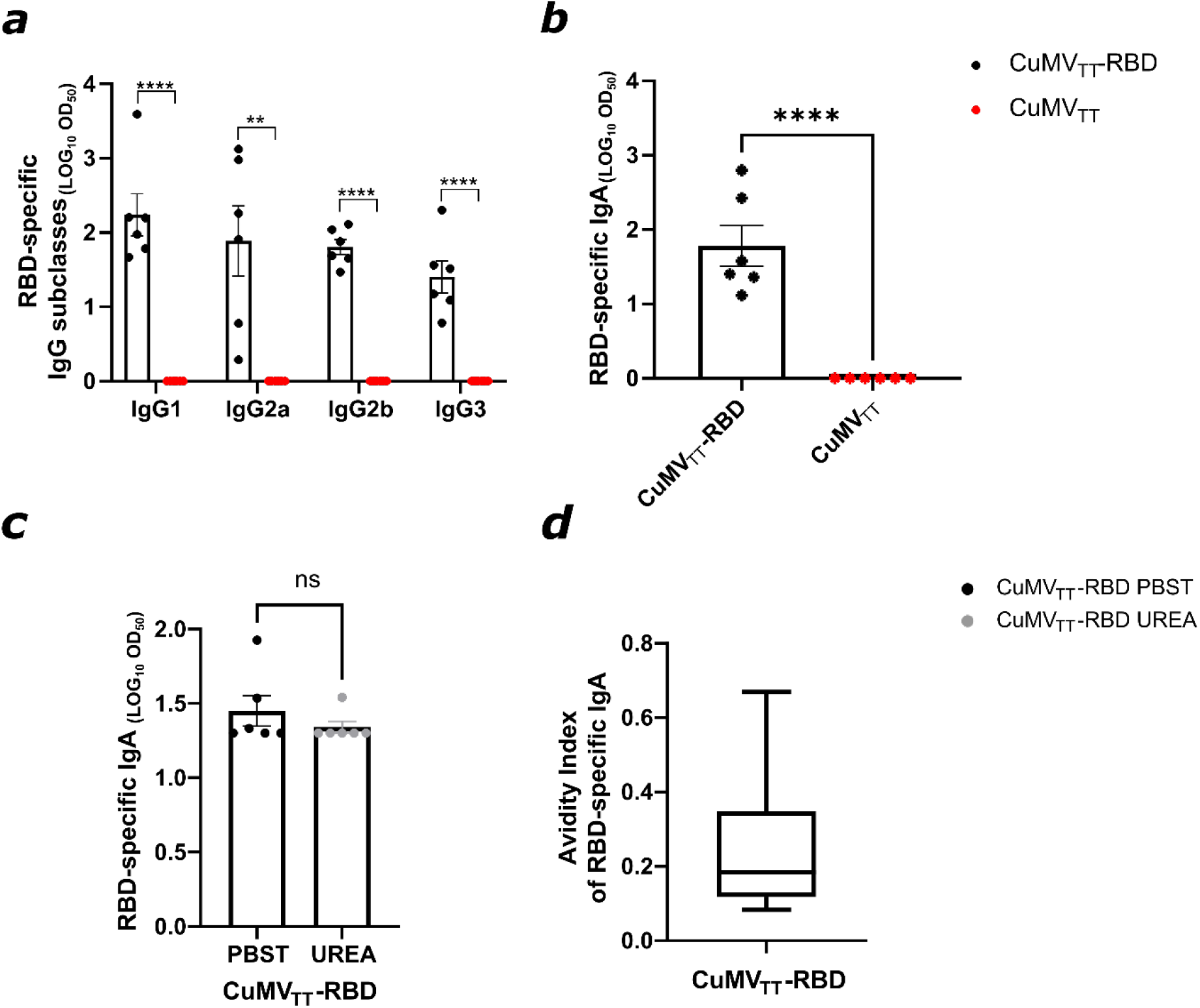
Intranasal immunization with CuMV_TT_-RBD promotes isotype switching to IgA and leads to balanced IgG subclass responses. a) RBD-specific IgG1, IgG2a, IgG2b and IgG3 titer for the groups vaccinated with CuMV_TT_-RBD vaccine or CuMV_TT_ control on day 42 measured by ELISA, LOG_10_ OD_50_ shown. b) RBD-specific IgA titer for the groups vaccinated with CuMV_TT_-RBD vaccine or CuMV_TT_ control on day 42 measured by ELISA, OD450 shown. c) RBD-specific IgA titer at day 42 from mice immunized with CuMV_TT_-RBD vaccine, LOG_10_ OD_50_ shown. Plates in duplicates: treated with PBSTween or 7M urea. d) Avidity index of RBD-specific IgA titer. Statistical analysis (mean ± SEM) using *Student’s t-test*. Control group *n=6*, vaccine group *n=6*. One representative of 2 similar experiments is shown.

### 3.4 CuMV_TT_-RBD induced immune sera is able to recognize VOCs

The mutation potential of SARS-CoV-2 is considered a burden in vaccine design and development, especially in terms of prolonged protection. Accordingly, we studied the capability of the induced RBD-specific IgG Abs in recognizing mutant RBDs of the different VOCs. Specifically, we have expressed and produced the following mutanted RBDs: RBD_K417N_, RBD_E484K_, RBD_N501Y_, RBD_K417N/E484K/N501Y_ and RBD_L452R/E484Q_^22^. Compared to RBD_wt_, RBD VoC specific IgG levels were slightly lower (Fig 4). However, the difference observed between RBD VOCs and RBD_wt_ IgG levels were statistically not significant (p=0.28). These findings indicate a broad potential protective capacity of Abs induced by i.n. vaccination with CuMV_TT_-RBD.

**Figure 4.**
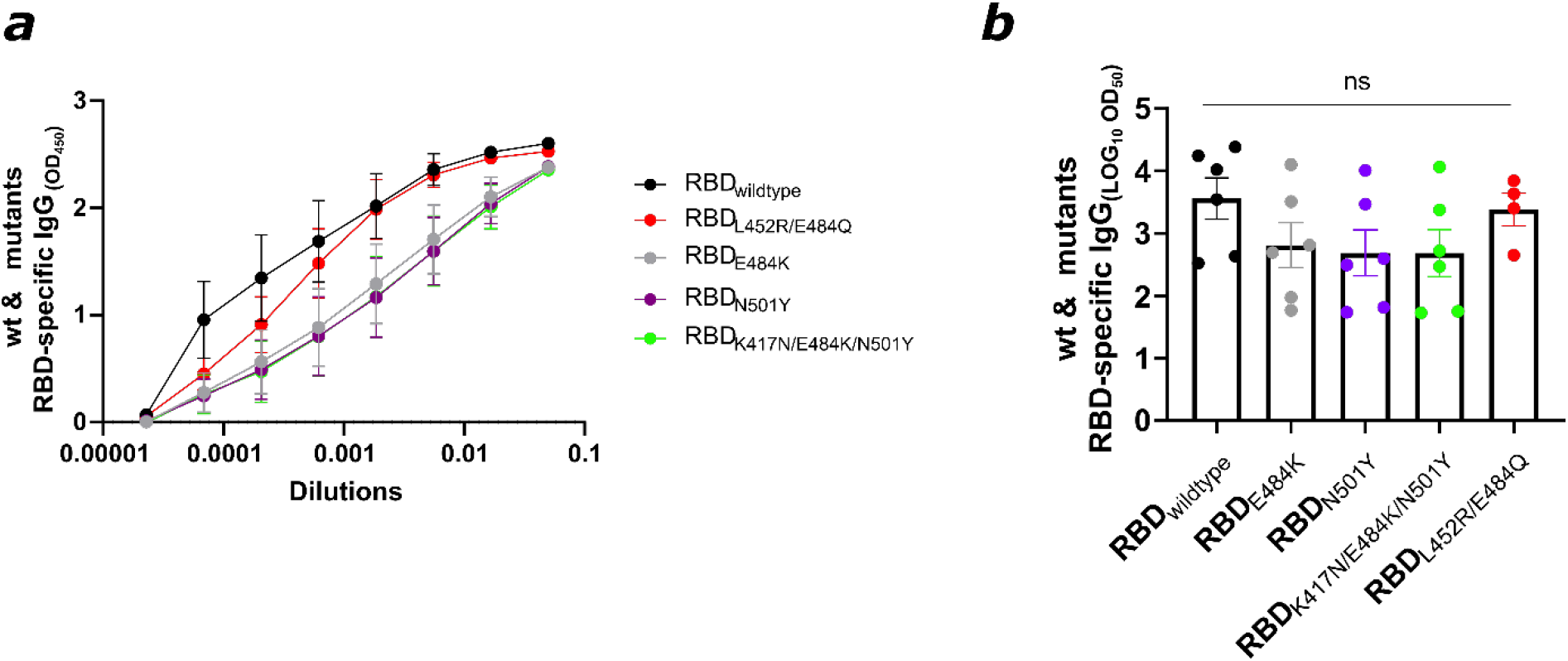
CuMV_TT_-RBD induced immune sera is able to recognize VOCs. RBD_wt_ and VOCs-specific IgG titers on day 49 for the group vaccinated with CuMV_TT_-RBD measured by ELISA, OD450 in a), LOG_10_ OD_50_ in b). Statistical analysis (mean ± SEM) using one-way ANOVA. Control group *n=6*, vaccine group *n=6*. One representative of 2 similar experiments is shown.

### 3.5 RBD- and spike-specific IgG and IgA were detected locally in BAL, with RBD-specific IgG2b dominating the local subclass response

To test the ability of the CuMV_TT_-RBD vaccine to induce a humoral immune response in the lung mucosa; BAL was collected two weeks after the booster injection (day 42) and assessed for RBD and spike protein specific IgG and IgA (Fig. 5a–f). RBD- and spike-specific IgG Abs were detected at equal levels in the BAL (Fig. 5a–c). However, IgA Abs in BAL were more abundant against RBD than against spike protein (Fig. 5d–f). We have also assessed the induced RBD-specific IgG subclasses in BAL. Interestingly, the local mucosal IgG response was less balanced than the serum response and dominated by IgG2b (Fig. 5g). Next, we tested the quality of the induced RBD-specific IgA in BAL samples, the results confirmed that about 60% of detected IgA Abs in BAL were of high avidity (Fig. 5h and 5i).

**Figure 5.**
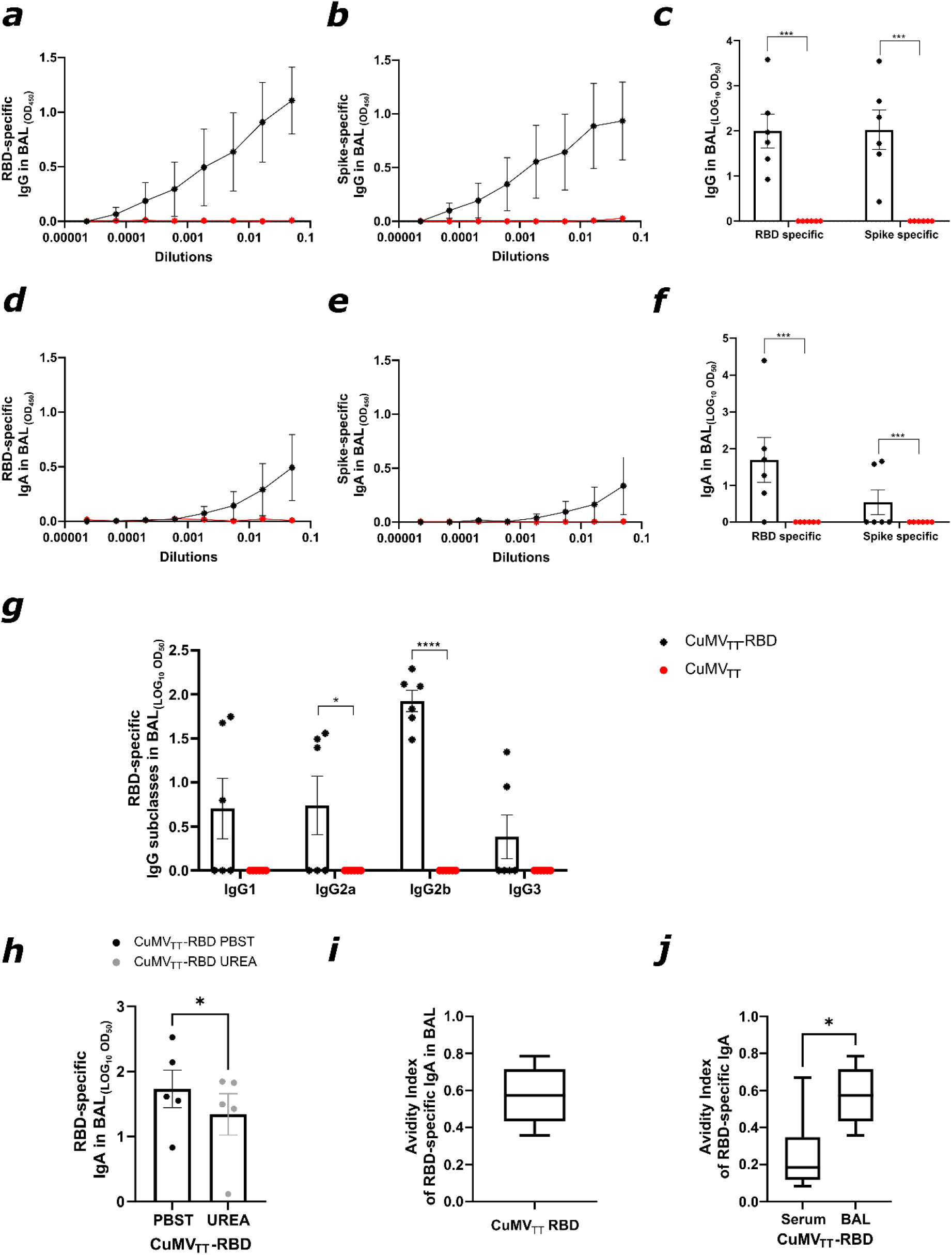
RBD- and spike-specific IgG and IgA were detected locally in BAL, with RBD-specific IgG2b dominating the local subclass response. a-c) RBD-(a,c) and spike-(b,c) specific IgG titer in BAL for the groups vaccinated with CuMV_TT_-RBD vaccine or CuMV_TT_ control, measured by ELISA, OD450 shown in a,b), LOG_10_ OD_50_ shown in c). d-f) RBD-(d,f) and spike-(e,f) specific IgA titer in BAL for the groups vaccinated with CuMV_TT_-RBD vaccine or CuMV_TT_ control, measured by ELISA, OD450 shown in d,e), LOG_10_ OD_50_ shown in f). g) RBD-specific IgG1, IgG2a, IgG2b and IgG3 titer in BAL for the groups vaccinated with CuMV_TT_-RBD vaccine or CuMV_TT_ control, measured by ELISA, LOG_10_ OD_50_ shown. h) RBD-specific IgA titer in BAL from mice immunized with CuMV_TT_-RBD vaccine, LOG_10_ OD_50_ shown. Plates in duplicates: treated with PBSTween or 7M urea. i) Avidity index of RBD-specific IgA titer. j) Avidity indexes of RBD-specific IgA Abs found in BAL and serum after CuMV_TT_-RBD immunization. Statistical analysis (mean ± SEM) using *Student’s t-test*. Control group *n=6*, vaccine group *n=6*. One representative of 2 similar experiments is shown.

### 3.6 Intranasal administration of CuMV_TT_-RBD induced RBD-specific IgG and IgA plasma cells locally and systemically

In order to characterize the humoral immune response upon i.n. CuMV_TT_-RBD vaccination, specific plasma blasts were quantified in lymphoid organs and lung tissue. To this end, spleen, BM and lung were collected on day 42 and analysed for the presence of RBD-specific IgG and IgA secreting plasma cells. As shown in Figure 6, IgG secreting plasma cells were detected in all investigated tissues. Around 25 IgG secreting plasma cells were found per two million cells seeded in spleen and BM. In lung, this ratio was ten-fold higher because the same amount of IgG plasma cells was observed while ten times less cells were seeded. IgA secreting cells were detected in all three tissues, however at a lower level compared to IgG secreting plasma cells. RBD-specific IgA producing plasma cells in lung were thereby about ten times more abundant compared to spleen or BM (Fig. 6a–c). In overall term, i.n. vaccination with CuMV_TT_ induced a systemic humoral immune response which was accompanied by a potent local humoral immune response in the lung.

**Figure 6.**
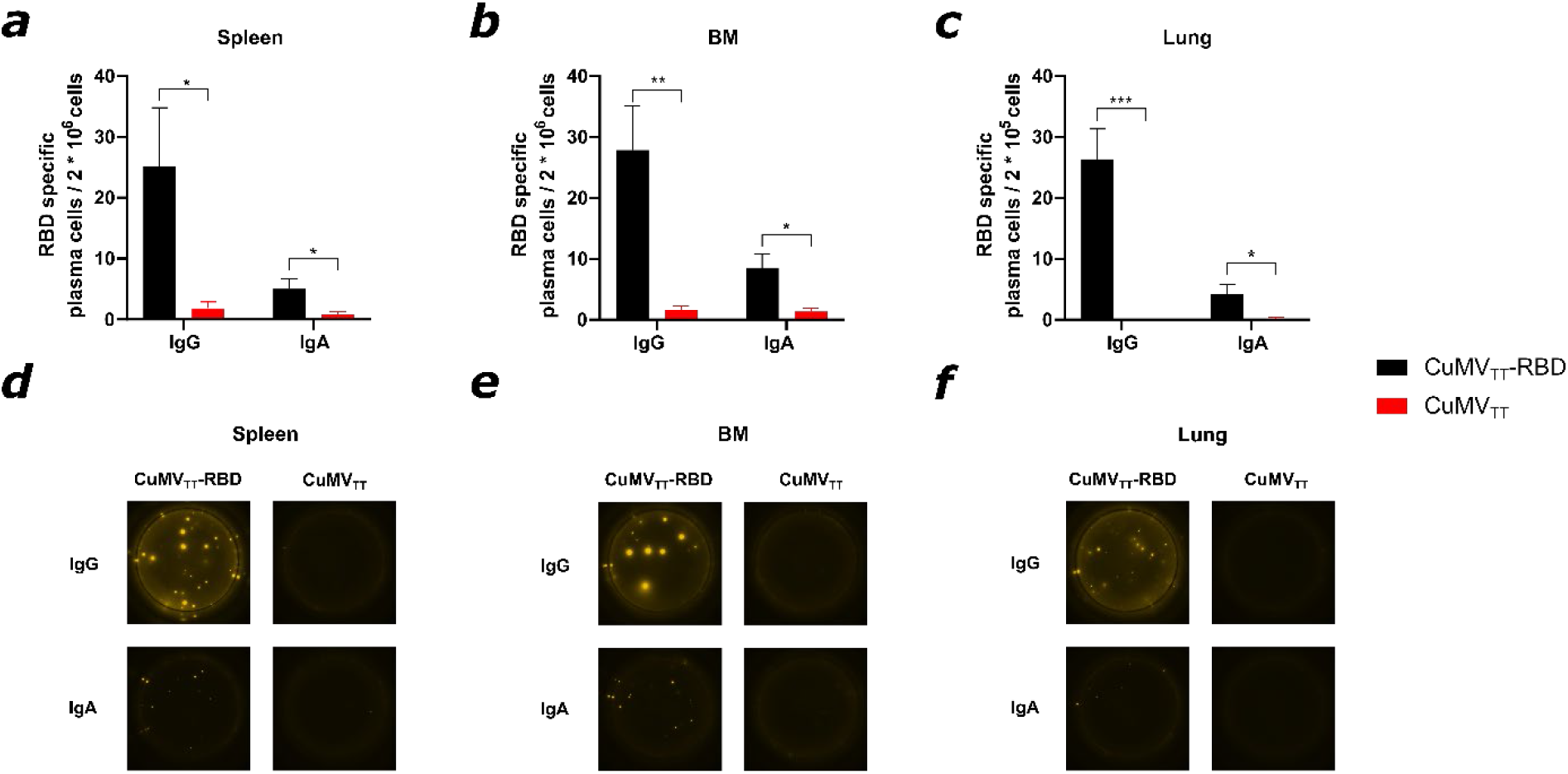
Intranasal administration of CuMV_TT_-RBD induced RBD-specific IgG and IgA plasma cells locally and systemically. a-c) Number of RBD specific IgG and IgA secreting cells per 2* 10^6^ seeded cells in spleen (a), BM (b) or per 2* 10^5^ seeded cells in lung (c) after immunization with CuMV_TT_-RBD vaccine or CuMV_TT_ control, detected by Fluorospot. d-f) Representative pictures of wells seeded with cells out of spleen (d), BM (e), lung (f). Statistical analysis (mean ± SEM) using *Student’s t-test*. Control group *n=6*, vaccine group *n=6*. One representative of 2 similar experiments is shown.

### 3.7 Abs induced by intranasal vaccination are capable of neutralizing SARS-CoV-2 and its variants

To test the ability of the sera of immunized mice to inhibit binding of RBD to ACE2, a biolayer interferometry competition assay was performed. Accordingly, RBD was immobilised onto anti-His biosensors and binding capacity of ACE2 to RBD in the presence of serum samples was quantified. As depicted in Figure 7a, the binding of ACE2 to RBD was reduced in the presence of sera from CuMV_TT_-RBD vaccinated mice. In contrast, no binding inhibition was observed in the presence of sera from CuMV_TT_ control mice. Percentage of ACE2 to RBD binding inhibition of individual mice is shown in Figure 7b. Interestingly, the binding inhibition correlated with RBD-specific IgG titers in serum (R value = 0.78) (Fig. 7c), indicating higher RBD specific IgG titre in serum are more efficient at blocking RBD-ACE2 interaction.

**Figure 7.**
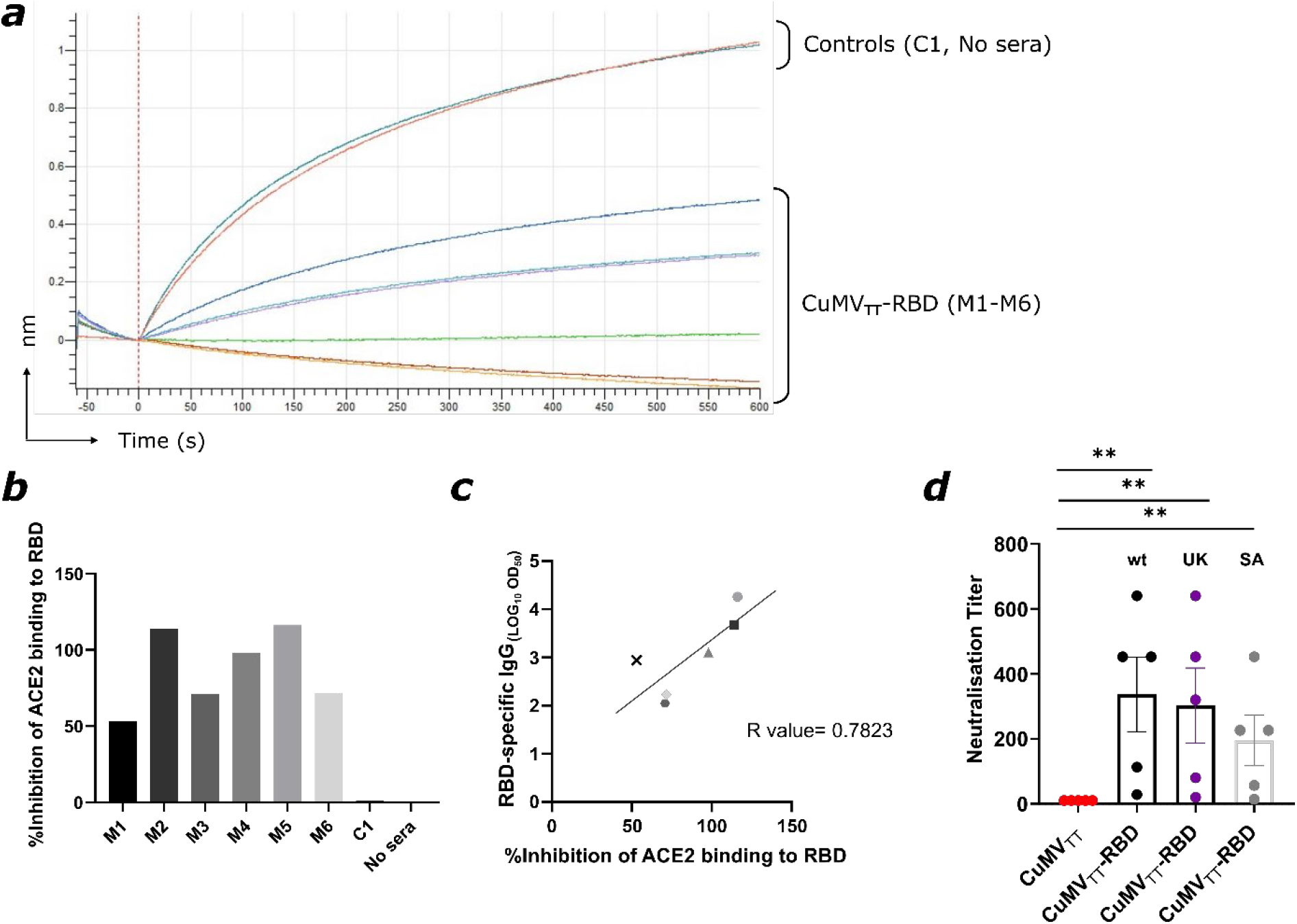
Abs induced by intranasal vaccination are capable of neutralizing SARS-CoV-2 and its variants. a) BLI-evaluation of ACE2 binding to RBD in the presence of vaccinated mice sera or controls. Percentage of ACE2 to RBD binding inhibition of the individual mice vaccinated with CuMV_TT_-RBD, control mouse vaccinated with CuMV_TT_ (C1) and with no serum. c) Correlation of RBD-specific LOG_10_ OD_50_ IgG titer with the percentage of ACE2 to RBD binding inhibition from individual mice sera. Statistical analysis (mean ± SEM) using Pearson Correlation test. d) Neutralization titer againstSARS-CoV-2 wt, SA and UK isolates. Statistical analysis (mean ± SEM) using *Student’s t-test*. Control group *n=6*, vaccine group *n=6*. One representative of 2 similar experiments is shown.

For the evaluation of viral neutralization capacity, sera from CuMV_TT_-RBD and CuMV_TT_ immunized mice were tested in a CPE-based neutralization assay. To this end, sera from immunized mice were assessed for their ability to prevent cytopathic effects of wt SARS-CoV-2 as well as VOCs on Vero cells *in vitro*. As shown in Figure 7d, all sera from CuMV_TT_-RBD vaccinated mice were able to neutralize the wt SARS-CoV-2 as well as SA and UK variants with high neutralization titers reaching to (1:600). In contrast, no neutralizing capacity was determined for sera from CuMV_TT_ immunized mice.

## 3. Discussion

The immune responses of the mucosal compartments are considered an early and essential line of defense against harmful pathogens such as SARS-CoV-2^23^. The majority of mucosal vaccines have been administered via oral or nasal routes, with the rectal, ocular, sublingual or vaginal routes being less often used^24^. Ideally, an effective mucosal vaccine would induce both local and systemic responses including the distant mucosal tissues. Accordingly, i.n. administration of a vaccine appears to be a promising strategy.

Seven ongoing clinical trials are currently testing the efficacy of i.n. vaccination against COVID-19 and are based on live-attenuated virus, viral-vectors or protein subunits. Some drawbacks from using attenuated viruses or viral-vectors includes: pre-existing Abs that can impair the vaccine efficacy and the risk of reversion of the live-attenuated viruses specially in newborns or immunocompromised people. In the current study, we have tested the immunogenicity and efficacy of a conventional vaccine based on VLPs displaying RBD of SARS-CoV-2 for i.n. administration. The multi-protein VLPs platform does not contain any genetic materials for replication and thus are considered a safe platform for vaccine development. Marketed vaccines against human-papilloma virus (HPV), hepatitis-B (HBV), hepatitis-E virus (HEV) and malaria are based on VLPs^19,25^. Our immunologically optimized CuMV_TT_-VLPs incorporate a universal T_H_ cell epitope derived from tetanus toxin (TT) and are self-adjuvanted with prokaryotic ssRNA, a potentTLR 7/8 agonist. We have shown in previous studies the essential role of TLR-7 signaling in licensing the generation of secondary plasma cells as well as the production of systemic IgA Abs. Such processes are usually dependent on TLR7-expression in B cells; however, in contrast to systemic IgA, a successful induction of mucosal IgA requires TLR signaling in dendritic cells (DCs)^26,27^. Our results here show effective induction of a systemic response of RBD-specific IgGs one week after the priming dose which increased significantly (*p*<0.001) following the booster dose on day 28 and antibodies showed a high degree avidity maturation. Moreover, a significant increase in RBD-specific IgA in serum was observed. Along with this, RBD-specific IgA and IgG secreting plasma cells were detected in spleen, BM and within lung tissues. Especially, IgG secreting cells showed a high Ab secretion rate. Consequently, i.n. vaccination with CuMV_TT_-RBD is able to induce a strong systemic humoral immune response.

As COVID-19 presents a respiratory disease and the virus is invading through the respiratory system, an enhanced immunological local protection in the lung should be in the focus when it comes to vaccine design. However, all vaccines currently licensed are applied intramuscularly^8^, thus ignoring this aspect. By applying CuMV_TT_-RBD i.n. instead of subcutaneously^28^ we were able to induce RBD- as well as spike-specific IgA and IgG Abs in the lung. IgA localized in lung mucosa has previously been shown to be of major importance for SARS-CoV-2 neutralization^29^. Furthermore, IgA may neutralize the virus in the lung without causing inflammation^10^. The fact that, around 60% of the RBD specific IgA Abs in the lung were of high avidity confirms the quality of the local humoral immune response upon intranasal vaccination with CuMV_TT_-RBD. Interestingly, only 20% of the RBD specific IgA Abs were of high avidity in serum. This significant difference might be explained by the 2 different forms of IgA: While IgA Abs at mucosal sides are mostly found in a dimeric form, serum IgA is usually present monomeric form^30^. In addition to IgA, IgG in the lung might also mediate protection^9^. By passive transudation across alveolar epithelium, IgG can pass from blood into the lower lung and from there by the mucociliary escalator further be carried to the upper respiratory tract and nasal passages. However, only at high serum concentrations local protection through IgG is achieved in the lung^9^.

While RBD-specific IgG subclass response in serum was well balanced, IgG2b was the most abundant subclass in BAL. Besides IgG2a, IgG2b is the only subclass that binds all three activating Fc receptors (FcγRI, FcγRIII, FcγRIV) and the only inhibitory receptor (FcγRIIB) in mice^31^. IgG2b Abs therefore mediate a wide variety of effector functions, which is of key importance in the maintenance of immune protection.

SARS-CoV-2 has the ability to mutate, albeit a proof-reading system is in place to keep the large genome of almost 30 kD genetically stable^32^. Previously, we have described that a single N501Y mutation increased the binding affinity to ACE2, but could still be detected by convalescent sera. Contrary were the results with the E484K mutation, where no enhanced binding to ACE2 was shown but much lower recognition by convalescent sera. Triple mutant RBD (K417N/E484K/N501Y) exhibited both features: stronger affinity to ACE2 and much lower detection by convalescent sera^33^. Since vaccines optimally mediate protection for many years, vaccine induced Abs should therefore be able to recognize new virus variants as well. In the present study we could show that i.n. applied CuMV_TT_-RBD induced serum IgG Abs that are able to recognize wt RBD as well as numerous RBD VOCs.

A crucial milestone in vaccine development is effective neutralization of the virus. Sera induced after i.n. administration of CuMV_TT_-RBD could completely inhibit the cytopathic effect of wt SARS-CoV-2 as well as other VOCs, specifically SA and UK variants. This may be explained by the highly repetitive, rigid antigenic surface array of VLPs which are spaced by 5 nm and displaying RBD domains at a spacing of 5-10nm ^34^. Such array of highly organized epitopes is considered a pathogen-associated structural patterns (PASPs) which are recognized by the immune system (the Nature Rev Immunol paper). In contrast, naturally induced Abs by SARS-CoV-2 are low in number and wane rapidly^35^.

It has been shown that B and T cells are primed by mucosal vaccination or natural infection, express receptors which promote homing of these cells to mucosal sites as Ab-secreting cells or effector or tissue-resident T cells^36^. We have shown in our previous studies that VLP-specific T_H_ cell response mediate specific B cell isotype-switch. Furthermore, the packaged RNA in VLPs can drive CD8 as well as T_H_1 responses^37^. Investigating the role of T cells after i.n. vaccination using VLPs is an area we are currently investigating.

Collectively, we have shown in this study that our COVID-19 vaccine candidate CuMV_TT_-RBD is highly immunogenic and capable of inducing both mucosal and systemic IgG and IgA response against SARS-CoV-2 upon i.n. administration. The induced Abs could effectively recognize and neutralize wt as well as the emerging VOCs. The ability of the vaccine candidate to stop nasal viral shedding and transmission is currently under investigation. As COVID-19 pandemic continues to present a global threat to human health, it seems rational to further develop an i.n. vaccine based on conventional platform.

## 4. Methods

### 4.1 Mice

All *in vivo* experiments were performed using (8-12 weeks old) wild-type (wt) female BALB/cOlaHsd mice purchased from Harlan. All animal procedures were conducted in accordance with the Swiss Animal Act (455.109.1-5 September 2008) of University of Bern. All animals were treated for experimentation according to protocols approved by the Swiss Federal Veterinary Office.

### 4.2 Protein expression and purification

RBD_wt_ of SARS-CoV-2 and mutant RBDs (RBD_K417N_, RBD_E484K_, RBD_N501Y_, RBD_K417N/E484K/N501Y_, and RBD_L452R/E484Q_) were expressed using Expi293F cells (Gibco, Thermo Fisher Scientific, Waltham, MA, USA). The amino acid (a.a.) sequence of each RBD was inserted into a pTWIST-CMV-BetaGlobin-WPRE-Neo vector (Twist Bioscience, San Fransico, CA, USA). RBD-His Tag construct was further transformed into competent XL-1 Blue bacterial cells. After plasmid purification, 50μg of the plasmid was then transfected into Expi293F cells at a density of 3×10^6^ cells/mL in a 250mL shaking flask using the ExpiFectamine 293 Transfection Kit (Gibco, Thermo Fisher Scientific, Waltham, MA, USA). Ninety-six hours later, the supernatant containing RBD was harvested and dialyzed with PBS. RBD protein was then captured using His-Trap HP column (GE Healthcare, Wauwatosa, WI, USA) or HiTrap TALON crude column (Cytiva, Uppsala, Sweden). Fractions were collected and concentrated. Buffer-exchanged to PBS was carried on using Vivaspin 20 5KDMWCO spin column (Sartorius Stedim Switzerland AG, Tagelswangen, Switzerland). Human ACE2 protein His Tag and SARS-CoV-2 spike were purchased from Sino Biological, Beijing, China.

### 4.3 CuMV_TT_-VLP expression and production

Expression and production of CuMV_TT_ VLP was described in detail in Zeltins et al.^15^.

### 4.4 Development of CuMV_TT_-RBD vaccine

RBD_wt_ was conjugated to CuMV_TT_ using the cross-linker succinimidyl 6-(beta-maleimidopropionamido) hexanoate (SMPH) (Thermo Fisher Scientific, Waltham, MA, USA) at 7.5 molar excess to CuMV_TT_ for 30 min at 25 °C. The coupling reactions were performed with molar ratio RBD/CuMV_TT_ (1:1) by shaking at 25 °C for 3h at 1200 rpm on a DSG Titertek (Flow Laboratories, Irvine, UK). Unreacted SMPH and RBD proteins were removed using Amicon Ultra 0.5, 100 K (Merck Millipore, Burlington, MA, USA). VLP’ samples were centrifuged for 2 min at 14,000 rpm for measurement on ND-1000. Coupling efficiency was calculated by densitometry (as previously described for the IL17A-CuMV_TT_ vaccine [28]), with a result of approximately 20% to 30% efficiency.

### 4.5 Electron microscopy

Physical stability and integrity of the candidate vaccine CuMV_TT_-RBD were visualized by transmission electron microscopy (Philips CM12 EM). For imaging, sample-grids were glow discharged and 10μL of purified CuMV_TT_-RBD (1.1mg/mL) was added for 30s. Grids were washed 3x with ddH_2_O and negatively stained with 5 μL of 5% uranyl acetate for 30s. Excess uranyle acetate was removed by pipetting and the grids were air dried for 10min. Images were taken with 84,000x and 110,000x magnification.

### 4.6 Vaccination regimen

Wild-type BALB/cOlaHsd mice (8-12 weeks, Harlan) were vaccinated intranasally (i.n.) with 40μg with either CuMV_TT_-RBD vaccine or CuMV_TT_ VLPs as a control in a volume of 40μL without any adjuvants (20μg in each nostril). The mice were boosted with an equal dose at day 28 and bled until day 49. Serum was collected on a weekly basis via tail bleeding and the serum was isolated using Microtainer Tube (BD Biosciences, USA). For Fluorospot samples and Bronchoalveolar lavage (BAL), wt BALB/cOlaHsd mice (8-12 weeks, Harlan) were vaccinated i.n. as indicated above. Serum samples were collected on day 35.

### 4.8 Enzyme-Linked Immunosorbent Assay (ELISA)

To determine the total IgG Abs against the candidate vaccine CuMV_TT_-RBM in sera of vaccinated mice, ELISA plates were coated with SARS-CoV-2 RBD_wt_ or spike protein (Sinobiological, Beijing, China) at concentrations of 1μg/mL overnight. ELISA plates were washed with PBS-0.01% Tween and blocked using 100μL PBS-Casein 0.15% for 2h at RT. Sera from vaccinated mice serially diluted 1:3 starting with a dilution of 1:20 and incubated for 1h at RT. After washing with PBS-0.01%Tween, goat anti-mouse IgG conjugated to Horseradish Peroxidase (HRP) (Jackson ImmunoResearch, West Grove, Pennsylvania) was added at (1:2000) and incubated for 1h at RT. ELISA was developed with tetramethylbenzidine (TMB), stopped by adding equal 1M H_2_SO_4_ solution, and read at OD_450_ nm or expressed as Log_10_ OD_50_. Detecting RBD-specific IgGs against mutated RBDs was carried out in a similar way.

To assess the subclass Ab response, the same procedure was performed. The following secondary Abs have been used: Rat anti-mouse IgG1 (BD Pharmingen, Cat. 559626, 1:2000 dilution), biotinylated mouse anti-mouse IgG2a (Clone R19-15, BD Biosciences, Cat No 553391, USA, 1:2000 dilution), goat anti-mouse IgG2b (Invitrogen, Ref. M32407, 1:2000 dilution), goat anti-mouse IgG3 (Southern BioTech, Cat No 1101-05, 1:2000 dilution).

To detect IgA Abs the plates were coated with 1μg/mL RBD protein and goat anti-mouse IgA POX (ICN 55549, ID 91, 1:1000 dilution) was used as a secondary Ab. IgG depletion was performed prior to serum incubation. 10μL of Protein G beads (Invitrogen, USA) were transferred into a tube and placed into a magnet. The liquid was removed and 75.6μL diluted sera in PBS-Casein 0.15% was added to the beads and mixed. The tube was incubated on a rotator at RT for 10 minutes. The tubes were placed back into the magnet and ELISA was carried out as described above.

### 4.9 Avidity (ELISA)

To test the avidity of IgG and IgA Ab against RBD, the above-described protocol was expanded by an additional step. Following serum incubation at RT for 1h, the plates were washed once in PBS/0.01% Tween, and then washed 3x with 7M urea in PBS-0.05%Tween or with PBS-0.05% Tween for 5 min. After washing with PBS-0.05%Tween, goat anti-mouse IgG conjugated to Horseradish Peroxidase (HRP) (Jackson ImmunoResearch, West Grove, Pennsylvania) was added (1:2000) and incubated for 1h at RT. IgA Abs were detected by using a goat anti-mouse IgA POX (ICN 55549, ID 91, 1:1000 dilution) detecting Ab. Plates were developed with TMB as described above and read at OD_450_ nm.

### 4.10 Bronchoalveolar Lavage (BAL)

BAL samples were collected as described in Sun et al.^16^.

### 4.11 Isolation of Lymphocytes

**From lung samples:** lungs were perfused with 10mL of 1mM EDTA in PBS via the right ventricle of the heart to remove blood cells from the lung vasculature. Lungs were dissected and digested for 30min at 37°C using RPMI media (2% FBS+ Pen/Strep, glutamine, 10mM HEPES) containing 0.5mg/mL Collagenase D (Roche). The digested fragments were passed through a 70μm cell strainer (Greiner bio-one, Art. Nr. 542070) and RBCs were lysed using ACK buffer. Lymphocytes were isolated using 35% Percoll gradient.

**From spleen samples:** the spleen was collected from mice and transferred into 5mL RPMI media (2% FBS+ Pen/Strep, glutamine, 10mM HEPES). A single cell suspension was prepared by passing the spleen through a 70μm cell strainer. The suspension was collected and transferred into a falcon tube. The tube was centrifuged for 8 min at 4°C and 300 x g. ACK lysis was performed, media added and centrifuged for 8 min at 4°C and 300 x g. The pellet was resuspended in media.

**From bone-marrow (BM):** tibia and femur were collected from mice and transferred into 5mL RPMI media (2% FBS+ Pen/Strep, glutamine, 10mM HEPES). The BM cells were isolated using a syringe by rinsing the bones to flush out the cells. A single cell suspension was prepared by passing the spleen through a 70μm cell strainer on a petri dish. The suspension was collected and transferred into a falcon tube. Petri dish was washed with 5mL media and also added to the falcon tube. The tube was centrifuged for 8 min at 4°C and 300 x g. ACK lysis was performed, media added and centrifuged for 8 min at 4°C and 300 x g. The pellet was resuspended in media.

### 2.14 Fluorospot

Fluorospot assay was performed according to manufacturer’s protocol (FluoroSpot-protocol.pdf (mabtech.com). Briefly, Fluorospot plate (Mabtech, Cat no. 3654-FL) was coated with 100μL RBD (50μg/mL) per well, 4°C overnight. The next day, plate was washed with PBS and blocked for 30 min at RT by addition of 200μL incubation medium (RPMI with 10% FBS, glutamine, pen/strep, 10mM HEPES) per well. 2×10^6^ cells from BM, spleen as well as 2×10^5^ cells from lung were seeded in 120μL medium per well. Plate was incubated at 37°C (5% CO_2_) for 20h. Cells were removed and the plate was washed with PBS. For detecting IgG secreting plasma cells, a goat anti-mouse IgG biotin primary Ab (SouthernBiotech, Cat no. 1030-08) at (1:1000) dilution in PBS-0.1% BSA was used. For detecting IgA secreting plasma cells, goat anti-mouse IgA biotin (Mabtech, Cat no. 3865-6-250) at (1:500) dilution in PBS-0.1% BSA was used. 100μL of Abs were added per well and plate incubated for 2h at RT. Afterwards plate was washed with PBS and Streptavidin-550 (Mabtech, Cat no. 3310-11-1000) diluted in PBS-0.1% BSA 1:200 was added, 100μL per well for 1h at RT. Plate was washed with PBS and Fluorescence enhancer-II (Mabtech, Cat no. 3641) was added, 50μL per well for 15 min at RT. The plate was flicked and dried at RT. Finally, plate was read at 550nm by using the Fluorospot reader (Mabtech IRIS).

### 4.15 BLI-based Assay

Antibody’ competitive binding activity was measured on an Octet RED96e (Fortébio) instrument which allows real-time analysis due to the shift in the wavelength of the reflected light. Anti-Penta-HIS (HIS1K, Lot 2006292, FortéBio) biosensors were first loaded into a biosensor microplate and pre-hydrated in BLI assay buffer (PBS, 0.1% BSA, 0.02% Tween 20) for 10 min. 96-well microplates were loaded with 200 mL per well. The tips were immobilized with 15μg/mL Sars-Cov-2 spike RBD containing a His-Tag (Sino Biological, USA). After, they were loaded with sera from mice, diluted 1:20 in BLI assay buffer. Next, association with 50nM of human receptor ACE2 (Sino Biological, USA) diluted in BLI assay buffer was measured. To regenerate the tips, two additional steps with regeneration buffer (0.1M glycine, pH 1.5) and neutralization buffer (BLI assay buffer) were performed.

### 4.16 SARS-COV-2 wt and Variant of Concern (VOC) live viruses

The SARS-CoV-2 2019-nCov/Italy-INMI1 clade V (Wuhan), the B.1.1.7 (UK VOC) named England/MIG457/2020, the B.1.351 (South Africa VOC) namedhCoV19/Netherlands/NoordHolland_10159/2021, next strain clade 20H were all purchsed from Europen Virus Archive (EVAg). SARS-CoV-2 strains were propagated in VERO E6 cells (ATCC—CRL 1586) in T175 Flasks using Dulbecco’s Modified Eagle’s - high glucose medium (DMEM) (Euroclone, Pero, Italy) supplemented with 2mM L-glutamine (Lonza, Milano, Italy), 100 units/mL penicillin-streptomycin (Lonza, Milano, Italy and 2% fetal bovine serum (FBS) (Euroclo, Pero, Italy). All viral growth and neutralization assay with SARS-CoV-2 live viruses were performed inside the VisMederi Bisecurity Level 3 laboratories.

### 4.17 Micro-Neutralization Assay cytopathic-effect-based (CPE)

The Micro-Neutralization assay was performed as previously reported by Manenti et al. 2020^17^. Briefly, 2-fold serial dilutions of heat-inactivated mice serum samples were mixed with an equal volume of viral solution containing between 25 TCID50 of SARS-CoV-2^18^. The serum-virus mixture was incubated for 1 h at 37 °C in a humidified atmosphere with 5% CO2. After the incubation time, 100 μL of the mixture at each dilution point was passed to a 96-well cell plate containing a sub-confluent VERO E6 (ATCC—CRL 1586) monolayer. The plates were incubated for 3 days (Wuhan strain) and for 4 days (B.1.1.7 and B.1.351) at 37 °C in a humidified atmosphere with 5% CO2. The day of the read-out each well was inspected by means of an inverted optical microscope to evaluate the percentage of cytophatic effect (CPE) developed in each well. The neutralization titre has been reported as the reciprocal of the highest dilution of serum able to inhibit and prevent at least in 50% of cells the CPE.

### 4.18 Statistical analysis

Data were analyzed and presented as (mean ± SEM) using *Student’s t-test* or *one-way ANOVA* as mentioned in the figure legend, with GraphPad PRISM 9. The value of *p*<0.05 was considered statistically significant (\**p*<0.05, \*\**p*<0.01, \*\*\**p*<0.001, \*\*\*\**p*<0.0001).

### 4.19 Supplementary Materials

**Supplementary Figure 1. CuMV_TT_-RBD induces local and systemic RBD-specific IgG and IgA producing cells.** a-c) Number of RBD-pecific IgG and IgA secreting plasma cells in spleen (a), BM (b) and lung (c). One representative of 3 similar experiments is shown.

## Data availability statement

The datasets generated during and/or analysed during the current study are available from the corresponding author on reasonable request.

## Acknowledgement

This publication was funded by Saiba AG, the Swiss National Science Foundation (SNF grants 31003A 149925 and 310030-179459) and Inselspital Bern and supported by the European Virus Archive goes Global (EVAg) project which has received founding from the European Union’s Horizon 2020 research and Innovation Programme under grant agreement No 653316.

## Author contributions

Design of experiments: MB, MM, PK, DR

Methodology: DR, PK, AN, IB, AV, XC, AM, FV, GR, SW, MM

Acquisition of data, interpretation and analysis of data: DR, PK, MM, MB, AV, AN, AM, FV, GR, EM, MV, AZ

Writing, revision and editing of manuscript: DR, PK, MM, MB Technical, material and tool support: EM, AZ

Study supervision: MB, MM, MV

All authors read and approved the final manuscript.

