## Supplementary Figure 1 for "Intranasal administration of a VLP-based vaccine against COVID-19 induces neutralizing antibodies against SARS-CoV-2 and Variants of Concerns"

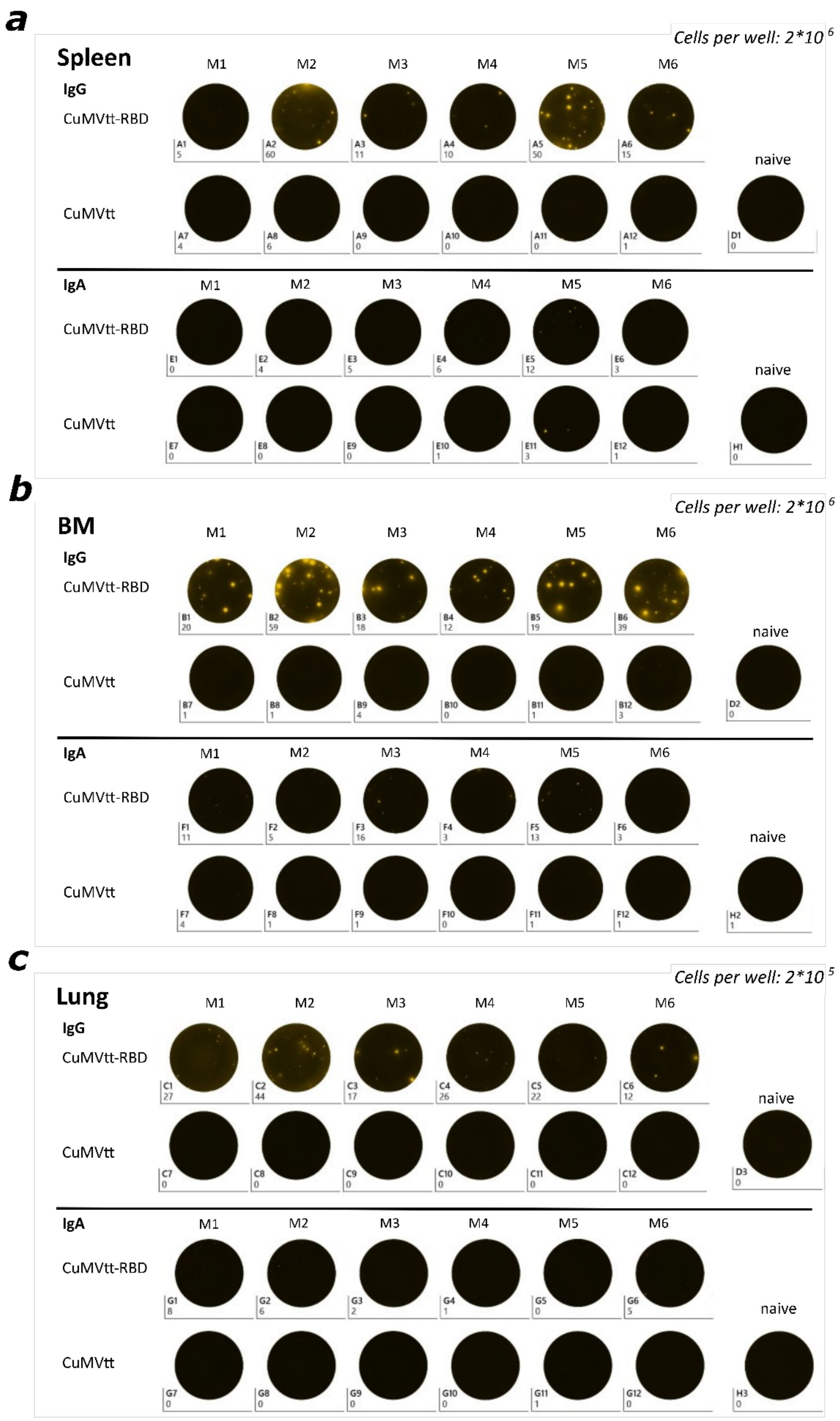


**Supplementary Figure 1. CuMV$_{TT}$-RBD induces local and systemic RBD-specific IgG and IgA producing cells.** a-c) Number of RBD -pecific IgG and IgA secreting plasma cells in spleen (a), BM (b) and lung (c). One representative of 3 similar experiments is shown.
